# Oxidative Stress Induced Senescence Gives Rise to Transcriptionally Distinct Physiological States

**DOI:** 10.1101/2022.05.18.492555

**Authors:** Nikolay Burnaevskiy, Junko Oshima, Alexander R. Mendenhall

## Abstract

As people age, the risk of cardiovascular disease, diabetes, cancer, and Alzheimer’s disease increases, making age itself the greatest risk factor for many human diseases. Thus, understanding the aging process can have profound consequences for human health. One striking feature of the aging process is the accumulation of senescent cells with age. When cells become damaged, they can enter a state of senescence which is a permanent cell cycle exit associated with the secretion of inflammatory cytokines. In mouse models of aging, the destruction of senescent cells with senolytic drugs delays age-associated decline and extends healthy lifespan. Yet, despite the wealth of accumulated knowledge, we do not entirely understand the biology of senescent cells. Prior work has shown that senescence is associated with increased variation in gene expression, suggesting that there may exist distinct transcriptional signatures of senescence. Understanding the different transcriptional physiological states of senescent cells would allow us to better treat them with cell-type-specific senolytic drugs. Here, we performed large-scale single-cell RNA-sequencing time series experiments to understand the development of transcriptional heterogeneity among senescent cell types. Our approach allowed us to observe and classify different transcriptional signatures of senescent cells as they emerged through time. We found that upon entering oxidative stress-induced senescence, separate subpopulations of cells were reproducibly adopting two distinct transcriptional states, one of which was associated with stress response and the second one with tissue remodeling. Our data suggest that a combination of senolytic drugs may be needed to more effectively eliminate senescent cells by targeting physiologically distinct sub-populations.

## Introduction

Aging incurs a tremendous socioeconomic cost. Life expectancy has significantly increased since the 20^th^ century. People spend a smaller portion of their life working, while concurrently requiring more resources for health care with increasing age. Age itself is the biggest risk factor for many diseases, including heart disease, cancer, and neurodegenerative diseases like Alzheimer’s Disease(Niccoli and Partridge, 2012). Thus, understanding the aging process and how to slow it, is a means of reducing the incidence of chronic diseases and decreasing the socioeconomic burden of aging.

Recent advances in aging research have shown excessive accumulation of senescent cells with age is a major factor that precipitates age-related physiological decline and mortality(Baker et al., 2016; Lopez-Otin et al., 2013; Xu et al., 2018) Senescence is a physiological state cells can enter in response to severe molecular damage; it is one of an array of natural cellular stress responses (Childs et al., 2017). To this end, the cell fate decisions vary widely depending on the cell type, intensity and duration of the stress stimuli: 1) repair the damage and resume normal function, 2) undergo apoptotic or necrotic cell death, 3) enter senescence, a state of stable proliferative arrest. More detailed characterization of senescent cells will help to better understand their normal physiological role and to develop novel treatments for selective clearance of excessively accumulated senescent cells (Childs et al., 2017). In the present study, we designed experiments to understand the senescence at single-cell level.

Senescent cell populations are heterogeneous. This is evidenced by senolytic compounds that work only on certain types of senescent cells (Zhu et al., 2015), and by transcriptionally distinct senescent cells produced by different types of molecular stress (Hernandez-Segura et al., 2017). Existence of different types or ‘flavors’ of senescent cells necessitates better characterization and maybe even better definition of senescence itself. Understanding how senescence manifests, and what types of senescent cell are there is paramount for understanding how to more effectively target them to improve the aging process.

The process of becoming a senescent cell itself may cause changes in physiological heterogeneity, in terms of cell-to-cell variation in gene expression. In some studies examining mouse and human cells, senescent cells had increased cell-to-cell variation in gene expression compared to their non-senescent counterparts (Bahar et al., 2006; Wiley et al., 2017). In nonsenescent cells, studies found aging is generally associated with increased cell-to-cell variation in gene expression (Cheung et al., 2018; Hernando-Herraez et al., 2019; Martinez-Jimenez et al., 2017; Salzer et al., 2018). Alternatively, at least one other recent study found that cell-to-cell variation in gene expression can also decrease with age (Kimmel et al., 2019). In any case, abnormal changes to heterogeneity also occur during aging (Mendenhall et al., 2021), and it is not entirely clear how they originate. Since, aging is accompanied by, and maybe even caused by, molecular damage, the cell-to-cell variation in gene expression among senescent and nonsenescent aged cells may have similar origins. It could be due to preexisting epigenetic differences, or due to random nature of molecular damage events. Understanding how and why cell-to-cell variation in gene expression changes with age may provide novel insights into the mechanisms of biological aging.

Here, we used large scale single-cell RNA-sequencing (scRNA-seq) to understand the mechanisms of cell-to-cell variation in gene expression upon induction of senescence. Changes in cell-to-cell variation in gene expression could be caused by subtypes of cells in distinct physiological states, or by purely stochastic noise in gene expression. In the first case, subgroups of cells would exhibit coordinated gene expression changes. In the latter case, individual genes would exhibit increased uncoordinated variance; there would be no coherent patterns. In order to distinguish between these possibilities, we performed transcriptome-wide analysis of gene expression at the single-cell level using the SPLITseq approach (Rosenberg et al., 2018). We performed time course experiments during senescence induction to generate whole-transcriptome data for thousands of individual cells across many individual experiments, finding reproducible patterns of gene expression variation. Our results indicate that cells after stress fall into distinct reproducible transcriptional clusters with different levels of metabolic signaling. This kind of reproducible, limited heterogeneity could be interpreted to be part of a functional bet-hedging mechanism in response to the same type of DNA damage or as a labor division between senescent cells, which may be required for complex processes involving senescent cells, such as wound healing.

## Results

### Two distinct transcriptional states emerge after oxidative stress

To establish senescent cells with oxidative stress, we exposed human IMR-90 fibroblasts to hydrogen peroxide for two hours (Chen et al., 2007). To compare senescence to other non-proliferative states, cells were cultured in low serum media (0.2% FBS) to induce quiescence or with nutlin-3a, an MDM-2 inhibitor (Figure 1a). Nutlin-3a was previously shown to induce either senescence or senescence-like arrest depending on conditions (Wiley et al., 2018). After one week, cells were stained with X-gal to analyze senescence-associated (SA) ß-galactosidase activity. We confirmed that both oxidative stress and nutlin-3a treatment induced robust SA ß-galactosidase staining (Figure 1b). In addition, cells after oxidative stress exhibited enlarged morphology, another classical marker of senescence (not shown). We collected cells from parallel plates (not used for X-gal staining) and processed them for single-cell RNA-sequencing (scRNA-seq) with SPLITseq 3’-tag protocol (Rosenberg et al., 2018). To determine the gene expression changes of stress induced senescence, we first compared average expression profiles of untreated cells and cells after oxidative stress. We found strong overlap with previously published signature of senescent fibroblasts (Hernandez-Segura et al., 2017): out of 1311 genes in the universal signature of senescent fibroblasts, 1072 were present in our dataset, and when adjusted for multiple comparison 205 of them had statistically significant change of expression in the same direction as in the universal signature (Tables S1).

**Figure 1.**
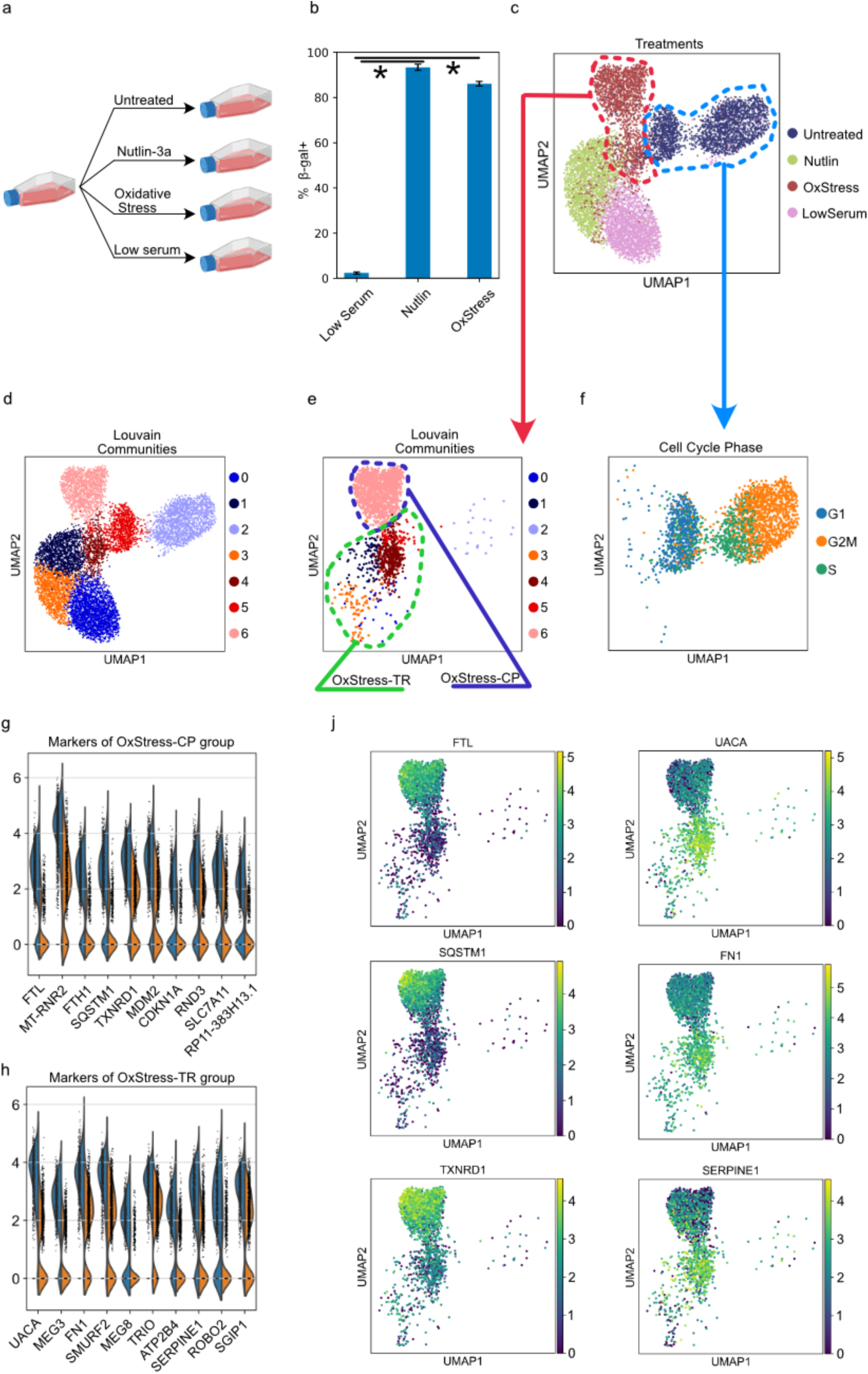
Two distinct senescence-like responses are observed upon oxidative stress. **a**. Experimental outline. Cells were treated with 2.5uM nutlin-3a to induce senescence/senescence-like arrest, or with 55uM H2O2 for 2 hours to induce senescence, or they were maintained for 7 days in 0.2% FBS media to induce quiescence. ‘Untreated’ cells were collected in the beginning of the experiment. **b**. Fraction of cells in each condition that were positive for senescence-associated ß-galactosidase as measured by X-gal staining. * indicates statistical significance with p<0.05 in t-test. **c**. Universal Manifold Approximation (UMAP)-based representation of transcriptional profiles of individual cells. Oxidative stress induces two distinct senescent states with different transcriptional signatures. **d**. Louvain community detection analysis of individual cells. **e**. Post-oxidative stress cells that were transcriptionally similar to nutlin-treated cells were labeled as OxStress-TR, and cells that were more transcriptionally distinct were labeled as OxStress-CP. **f.** Cell cycle phase of ‘Untreated’ cells were determined by scoring expression of cell cycle specific genes. **g, h**. Markers of OxStress-CP and OxStress-TR fractions were determined by Wilcoxon rank-sum test. Shown are top 10 statistically significant markers for each fraction. Also see Table S2. **j**. Expression of select markers of each fraction. Two OxStress fractions exhibit antagonistic pattern of expression of the markers.

To proceed to single-cell analysis, we performed dimensionality reduction and embedding with Universal Manifold Approximation (UMAP) (Becht et al., 2019). The results are shown in Figure 1c. Upon examining the UMAP space, we found that cells induced into senescence with oxidative stress fall into two distinct transcriptional clusters (Figure 1c, OxStress cells). This transcriptional heterogeneity was specific to post-oxidative stress cells, as both quiescent cells and nutlin-treated cells formed singular clusters (Figure 1c, ‘LowSerum’ and ‘Nutlin’ cells respectively). Louvain community detection analysis also indicated that two fractions of OxStress fall into distinct Louvain clusters, further underscoring their transcriptional heterogeneity (Figure 1d, e). Interestingly, one of the OxStress clusters localized closer to ‘Nutlin’ cells in the UMAP space, indicating stronger transcriptional similarity to nutlin-treated cells. The only other sample in our experiment that formed two clusters was the control sample with untreated cells which segregated based on their cell cycle phase (Figure 1c, f). We examined markers that differentiate two OxStress clusters from each other (Figure 1g, h, Table S2). Fraction of OxStress cells that was transcriptionally closer to nutlin-treated cells and quiescent cells was marked by expression of genes associated with cell motility/adhesion/organization: UACA, FN1, TRIO, SERPINE1, ROBO2. Hence, we labeled this fraction as ‘OxStress-TR’ (tissue remodeling) for further discussion (Figure 1e). Fraction of OxStress cells that was transcriptionally more distinct from quiescent and nutlin-treated cells was marked by expression of genes associated with cellular homeostasis (Figure 1g): FTL and FTH1 are subunits of ferritin, which stores intracellular iron, SQSTM1 is a regulator of autophagy, thioredoxin reductase TXRND1 and cystine/glutamate transporter SLC7A11 both help maintain cellular redox balance. Hence, this fraction of OxStress cells was labeled as ‘OxStress-CP’ (cytoprotective) (Figure 1e). As shown in Figure 1j, identified markers exhibit antagonistic expression pattern in ‘cytoprotective’ and ‘tissue remodeling. Thus, upon oxidative stress, senescent fibroblasts exhibited two distinct transcriptional signatures: cytoprotective response and response associated with tissue organization/remodeling.

It was important for us to distinguish technical noise associated with scRNAseq from genuine biological variation of gene expression (Mendelevich et al., 2021). Therefore, we performed independent repetitions of the experiment (Figure S1a-g). Most of the cells after oxidative stress were again positive for SA ß-galactosidase activity (Figure S1a). Consistent with the first experiment, we found that after oxidative stress, cells separated into two transcriptional fractions, while low serum- and nutlin-treated cells formed singular transcriptional communities (Figure S1b). Analysis of the two OxStress fractions confirmed that they are distinguished by the same reproducible markers (Figure S1e-g, Tables S3). Thus, the uncovered transcriptional heterogeneity was not a result of technical noise but was a reproducible biological heterogeneity.

We considered a possibility, that one of the OxStress fractions corresponds to cells that resumed proliferation after stress. However, proliferating cells in S and G2/M phases were clearly distinct from other clusters in transcriptional space (see Figures 1c, f) and only small number of cells from OxStress, LowSerum and Nutlin samples co-localized with untreated cells in S/G2/M phases in transcriptional space (see Figures 1c, f). Consistently, we observed large number of mitotic cells on untreated plates, but not on LowSerum-, OxStress-, or nutlin-treated plates (not shown). Hence, neither of the large transcriptional fractions of OxStress cells represent cells that resumed proliferation.

Next, we examined the possibility that either of the OxStress fractions may be representing cells that were somehow not strongly affected by oxidative stress. Two lines of evidence argue against this possibility. First, oxidative stress rendered a vast majority of cells SA ß-galactosidase-positive in the initial experiment, inconsistent with a possibility that a large fraction of OxStress cells were not strongly affected by oxidative damage (see Figure 1b). Next, we performed another independent experiment with few modifications: cells were treated with higher concentration of hydrogen peroxide and were treated with it twice (Figure S1 h-n). Almost 100% of cells were SA ß-galactosidase-positive after the treatment (Figure S1h), but the OxStress cells were still split into two transcriptional fractions (Figure S1i-k) distinguished by a similar set of marker genes as seen earlier (Figure S1i-n, Table S4). Thus, both fractions of OxStress cells represent SA ß-galactosidase-positive cells that were induced into senescence or senescence-like state by oxidative stress. In summary, our results show that, upon oxidative stress, most cells subsequently exhibit signs of senescence, and they fall into two distinct transcriptional states. The observed transcriptional heterogeneity is in line with recent studies showing changes in transcriptional heterogeneity *in vivo* and *in vitro* with aged cells (Bahar et al., 2006; Cheung et al., 2018; Hernando-Herraez et al., 2019; Martinez-Jimenez et al., 2017; Salzer et al., 2018; Wiley et al., 2017). Our results highlight that the term ‘senescent cells’ needs better, more precise or, on the opposite, a broader definition.

### Subgroups of oxidative stress-induced senescent cells exhibit distinct functional and metabolic signatures

To better characterize the observed fractions of senescent cells, we performed gene set enrichment analysis (GSEA) and compared these two fractions to each other and to cells from other conditions. We chose ‘Hallmarks’ gene sets that represent well-defined biological states or processes and reduce redundancy and noise (Liberzon et al., 2015). First, we compared OxStress-CP and OxStress-TR to quiescent (LowSerum-treated) cells and we noted that both fractions exhibited signs of stress. OxStress-CP cells expressed hallmarks ‘Unfolded Protein Response’, ‘Reactive Oxygen Species Pathway’, ‘p53 Pathway’ and ‘DNA Repair’, while OxStress-TR cells were enriched for a hallmark ‘Unfolded Protein Response’, but not others (Figure 2a, b). In a repetition experiment, both fractions were more enriched for hallmarks of stress when compared to quiescent cells, but OxStress-CP group was again enriched more (Figures S2a, b). Another, distinction between the two fractions was in the area of metabolism. OxStress-CP cells were again more distinct from quiescent cells and exhibited hallmarks of MTORC1 pathway (‘MTORC1 Signaling’, PI3K/AKT/MTOR Signaling’), ‘E2F Targets’, and ‘Oxidative Phosphorylation’. Hence, between the two fractions, OxStress-CP mounted broader stress response and seemed to be in a more active metabolic state compared to OxStress-TR cells.

**Figure 2.**
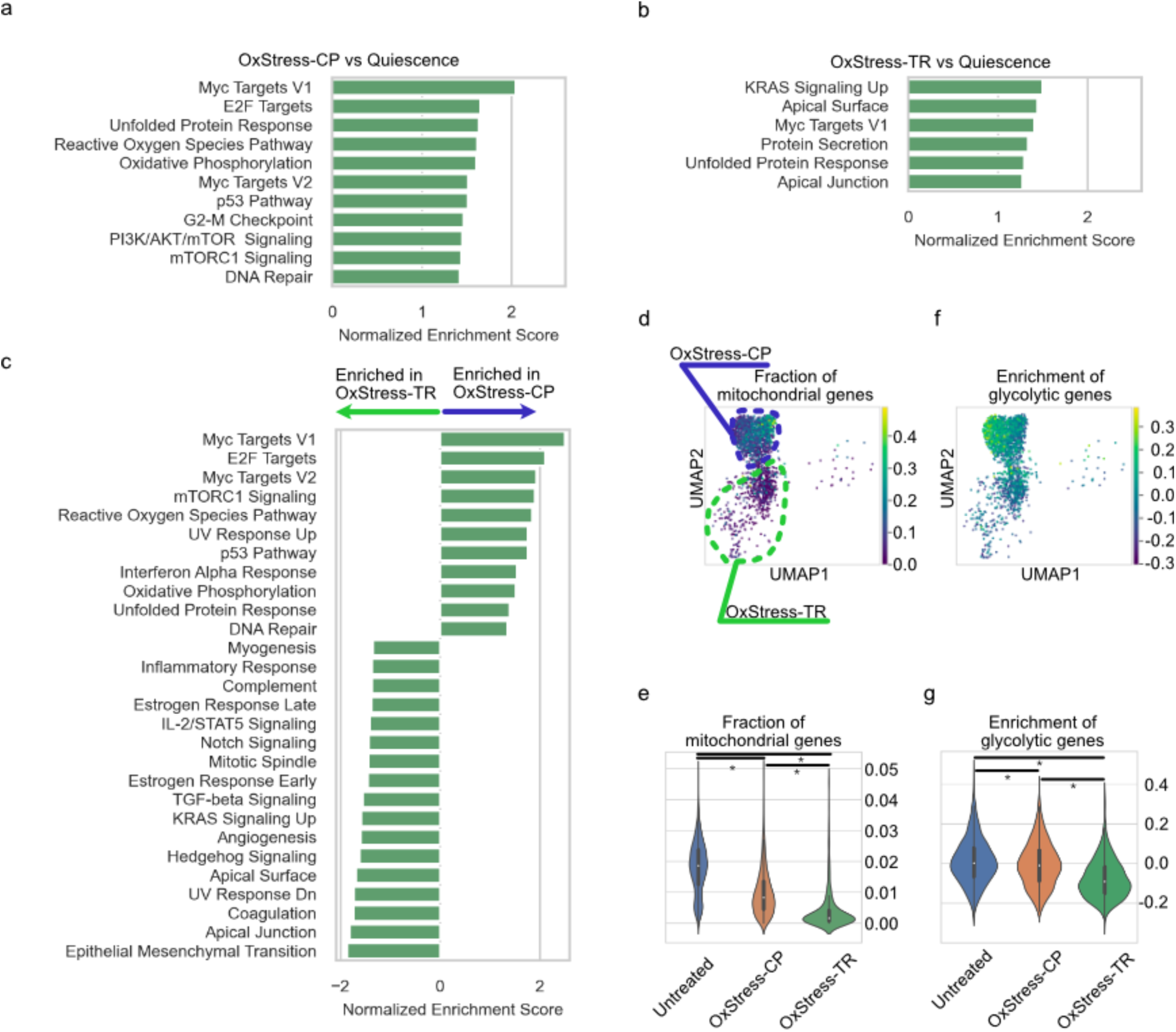
Gene Set Enrichment Analysis (GSEA) of OxStress fractions. **a**. ‘Hallmarks’ gene sets enriched in OxStress-CP compared to Quiescent cells (low serum-treated cells). **b**. Hallmarks gene sets enriched in OxStress-TR compared to Quiescent cells. **c**. Hallmarks gene sets enriched in OxStress-CP compared to OxStress-TR cells (positive values) or in OxStress-TR compared to OxStress-CP (negative values). **d**. Fraction of mitochondrial transcripts in transcriptome of individual OxStress cells. **e**. Fraction of mitochondrial transcripts in Untreated cells and OxStress-CP and OxStress-TR groups. **f.** Enrichment of glycolysis-related genes in individual OxStress cells. **g**. Enrichment of glycolysis-related genes in Untreated, OxStress-CP and OxStress-TR groups.

Next, we directly compared OxStress-CP and OxStress-TR fractions to each other (Figure 2c). Consistent with an earlier analysis, OxStress-CP cells were more metabolically active (hallmarks ‘Myc Targets V1’, ‘Myc Targets V2’, ‘E2F Targets’, ‘mTORC1 Signaling’, ‘Oxidative Phosphorylation’) and exhibited stronger activation of stress-related pathways (‘Reactive Oxygen Species’, ‘UV Response Up’, ‘p53 Pathway’, ‘Interferon Alpha Response’, ‘Unfolded Protein Response’, ‘DNA Repair’) than OxStress-TR cells. Also consistent with an earlier notion, OxStress-TR cells were more enriched for cell- and tissue-organization activities (‘Apical Surface’, ‘Apical Junction’, ‘Epithelial Mesenchymal Transition’ Hallmarks). Our analysis showed a similar difference between OxStress-CP and OxStress-TR fractions in a repetition experiment (Figure S2c). Hence oxidative stress induced senescence comprised two signaling states with distinct functional specializations.

We performed a more detailed analysis of the two OxStress fractions to understand how the transcriptional changes may relate to differences in cell physiology, focusing on mitochondrial and glycolytic metabolism. We found that two fractions differ in abundance of mitochondrial genes, with OxStress-CP cells being more enriched in mitochondrial transcripts (Figures 2d-e, S2d-e). However, when we compared OxStress cells to the untreated control, we found that the difference was not due to elevated expression of mitochondrial genes in OxStress-CP cells compared to non-stressed cells, but rather due to strongly decreased expression of mitochondrial genes in OxStress-TR cells compared to both OxStress-CP and untreated cells (Figure 2e, S2e). Importantly, abundance of mitochondrial transcripts was regressed out from gene expression datasets prior to initial analysis, so the appearance of the two post-stress cell fractions is not a technical artifact due to variable abundance of mitochondrial transcripts. We then examined expression of glycolytic enzymes as another measure of metabolic activity. OxStress-CP cells were again enriched in glycolytic transcripts, underscoring higher metabolic activity of this cell subset (Figures 2f-g, S2f-g). Interestingly, mitochondrial enrichment did not overlap with glycolytic enrichment within OxStress-CP subset in one experiment. Smaller size of OxStress-CP fraction in a repetition experiment did not allow such a detailed examination (Figure S2f). Thus, two fractions of post-stress cells differ in metabolic signaling, and both glycolytic and mitochondrial activity promote this difference.

### Transcriptional heterogeneity emerges early during stress response

We did further examination of the gene sets of the senescent cells fractions. We noticed that OxStress-TR fraction exhibited signatures of TGF-beta Pathway and Notch Signaling (Figures 2c, S2c). Remarkably, recent studies identified two senescence fates upon oncogene-induced senescence (OIS), one of which was characterized by inflammatory cytokines and the other one by Notch signaling (Hoare et al., 2016; Teo et al., 2019). Hence, we uncovered that these distinct fates arise not only upon OIS, but also upon stress induced senescence.

In the OIS model, TGF-beta-associated senescence (‘secondary senescence’) appeared to be a response to signals secreted by ‘primary’ senescent cells, where Notch was mediating a switch from ‘inflammatory’ fate to TGF-beta fate. We wanted to get better understanding of the senescence fate determination, and how it emerges. It is possible that all cells exhibit the same response to oxidative stress at early stages and the fate split happens later during stress response. Alternatively, heterogeneity may establish immediately after stress response. Discriminating between these possibilities will help better understand the mechanisms behind the distinct senescence states. To this end, we performed time course experiments in which cells were analyzed before stress (untreated control) as well as four hours, one day, two days, three days, four days, and seven days after stress (Figure 3a). Consistent with earlier experiments, cells at the final time point clustered into two transcriptional groups (Figure 3a). To better analyze emergence of distinct post-stress fates, we combined genes that reproducibly distinguish OxStress-CP and OxStress-TR fractions in all experiments into two gene sets (‘CP’ signature and ‘TR’ signature). We then scored all cells in a time course experiment to determine when ‘CP’ and ‘TR’ signatures emerge. Both signatures were the most pronounced at the final time point and were developing gradually over the course of time (Figure 3b, c). We then examined each time point more closely. Figure 3d shows cells and enrichment of the two signatures at the different time points. By following time course from the last time point to the first, we noticed that clear separation of the two fractions was not apparent until Day 7 (Figure 3d). At the earlier time points cells were mostly in a single transcriptional cluster starting from Day 1 (Figure 3d). Analysis was complicated at earlier time points because of the differences in cell cycle phases (Figure 3a, d). Notably, at all the time points between Day 1 and Day 7 there was clear antagonism between ‘CP’ and ‘TR’ signatures where cells were leaning toward one or another fate. Hence, we propose that although strong transcriptional separation establishes only at later time points, post-stress fate might be biased if not determined early after stress.

**Figure 3.**
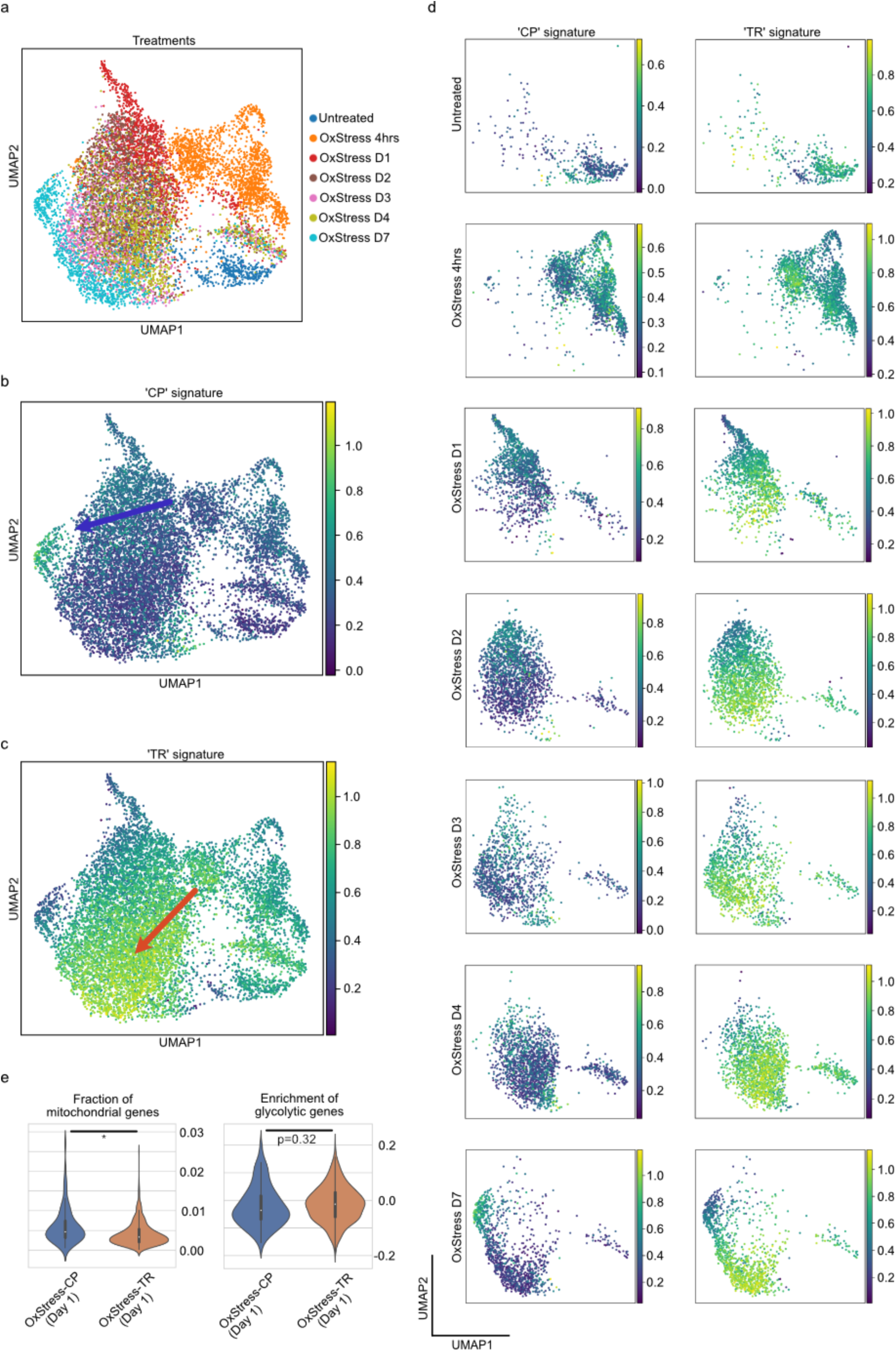
Cells exhibit early bias toward post-stress fates. Cells were collected at different time points after oxidative stress. **a**. UMAP representation of time course analysis. Cells are colored based on their sample of origin. **b**, **c**. Enrichment of type A and type B signatures across the time course. A and B signatures are sets of genes that distinguish type A and type B responses (see Table X). **d**. Enrichment of type A and type B signatures at individual time points. **e**. Fraction of mitochondrial transcripts and enrichment of glycolytic transcripts ‘A’ and ‘B’ group cells at Day 1. At this time point, cells were assigned to ‘A’ or ‘B’ depending on which respective signature was higher.

This bias we observed could have resulted from cell-to-cell differences existing before stress, from different levels of damage experienced by individual cells, or from interaction of these factors. In one repetition experiment cells separated into two transcriptional groups earlier during time course (Figures S3a-d), and another independent trial gave similar results to the first one (Figures S3f-j). In all independent trials, bias toward a particular post-stress fate was observed early during stress response. We then examined if metabolic difference between cell fractions also establishes early during stress response. We found that similarly to late time points, ‘TR’^high^ at Day 1 (cells in which ‘TR’ signature scored higher than ‘CP’ signature) exhibited lower abundance of mitochondrial transcripts compared to ‘CP’^high^ cells (Figure 3e, S3e,k), indicating metabolically less active state. Interestingly, abundance of glycolytic transcripts did not show such consistent difference between ‘CP’^high^ and ‘TR’^high^ cells at Day 1(Figure 3e, S3e,k). Thus, cell-to-cell difference in mitochondrial activity is among early signs of post-stress heterogeneity, while intercellular differences in glycolytic activity develops later. Overall, we observe early emergence of bias toward a particular senescence fate, CP or TR, suggesting that senescence fate is at least partially determined early during stress response.

Senescence is an alternate fate to apoptosis, which tends to occur when cells are severely damaged and retain the apoptosis capacity. Accordingly, we also determined if either of the OxStress fractions represented dying cells, e.g. apoptotic cells. We analyzed number of cells collected at each time point. Consistent with visual examination, the number of cells dropped between the untreated control and the four-hour time point (Figure S4). At the four-hour time point, we observed a large number of rounded, presumably dying cells (not shown). At later time points we observed neither ‘apoptotic’-like nor mitotic cells. Consistent with that, number of cells remain approximately the same at subsequent time points (Figure S4). While we cannot rule out that one of the fractions is destined for eventual cell death or on the opposite for eventual recovery and cell cycle re-entry, we found at least for a while, cells entering oxidative stress induced senescence represent a mix of at least two signaling states. This finding prompts further careful examination of senescent cells at single cell level for better understanding of senescence and identification of reliable markers of senescent cells.

### Increased stochastic noise is not a universal feature of senescence

After establishing that oxidative stress induced senescence is associated with variability of signaling states (signaling noise), we asked if senescence also increased stochastic noise, that is uncoordinated cell-to-cell variation of gene expression. Observable stochastic noise of gene expression strongly depends on expression level with lower expression being associated with increased noise due to both technical and biological factors (Elowitz et al., 2002; Mendelevich et al., 2021; Raser and O’Shea, 2004). Normalizing read counts across cells to the same value, which is a typical step in scRNAseq data analysis, can therefore give exaggerated noise estimates for samples with lower read counts. Hence, for this analysis we used non-normalized data. We calculated coefficients of variation (CV) as well as mean expression for each gene in OxStress (OxStress-CP and OxStress-TR combined) and untreated cells. Scatterplot at Figure 4a shows that expression variation for most genes is decreased in OxStress cells. The decrease is likely explained by increased mean expression level (Figure 4b). Indeed, we noticed that CV of some genes increased in OxStress cells relative to majority of other genes (shown in red in Figure 4a), and that coincided with decreased mean expression of these genes after oxidative stress (Figure 4b). To further examine the changes in intrinsic expression noise, we selected genes with similar average expression level in both post-stress and untreated conditions (Figure 4c). We found that CV of those genes changed little between two conditions (Figure 4d). To exclude effect of cell cycle, we also compared OxStress cells to quiescent (low serum-treated) cells and we got similar results (Figure 4e-h). Independent repetition produced the same results (Figure S5). Hence, we did not find evidence for increased stochastic noise upon oxidative stress induced senescence.

**Figure 4.**
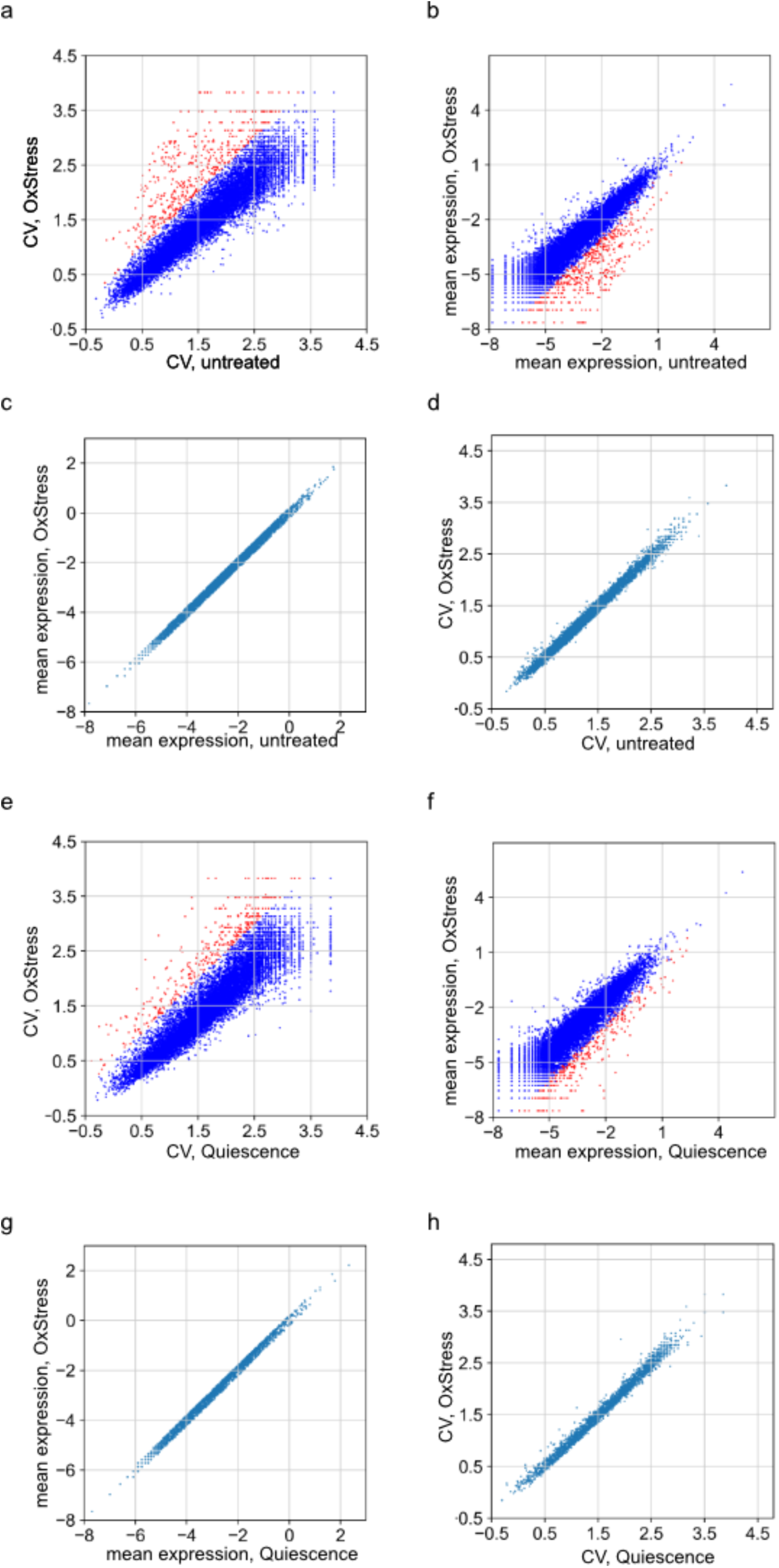
Increased stochastic noise of gene expression is not a universal feature of senescence. **a**. Coefficient of variation (CV) of genes in untreated and OxStress cells. **b**. Mean expression of genes in untreated and OxStress cells. **c**. Mean expression of genes with similar expression in untreated and OxStress cells. **d**. CV of genes with expression was similar in untreated and OxStress cells. **e**. Coefficient of variation (CV) of genes in quiescent and OxStress cells. **f.** Mean expression of genes in quiescent and OxStress cells. **g**. Mean expression of genes with similar expression in quiescent and OxStress cells. **h**. CV of genes with similar expression in quiescent and OxStress cells. All axes are in log scale.

### Expression of senescence markers in different fractions of post-stress cells

Identification of reliable markers of senescent cells for research and clinical use is an area of intense investigation. We used our single-cell data to examine expression of previously reported universal markers of senescent fibroblasts (Hernandez-Segura et al., 2017). When we cross-referenced fibroblasts’ universal senescence markers with our list of genes that distinguish OxStress-CP and OxStress-TR fractions (Tables X and X), we found genes that change expression stronger in one or another fraction. For some genes, average change of expression in OxStress cells was mostly driven by only one OxStress fraction (Figure 5). Some genes were mostly upregulated in OxStress-CP (Figure 5a) or in OxStress-TR cells (Figure 5b), and some were mostly downregulated in OxStress-CP (Figure 5c) or in OxStress-TR (Figure 5d). Hence, single cell analysis allows better characterization of senescence markers, since expression of certain markers may be mostly affected by a subset and not by all senescent cells.

**Figure 5.**
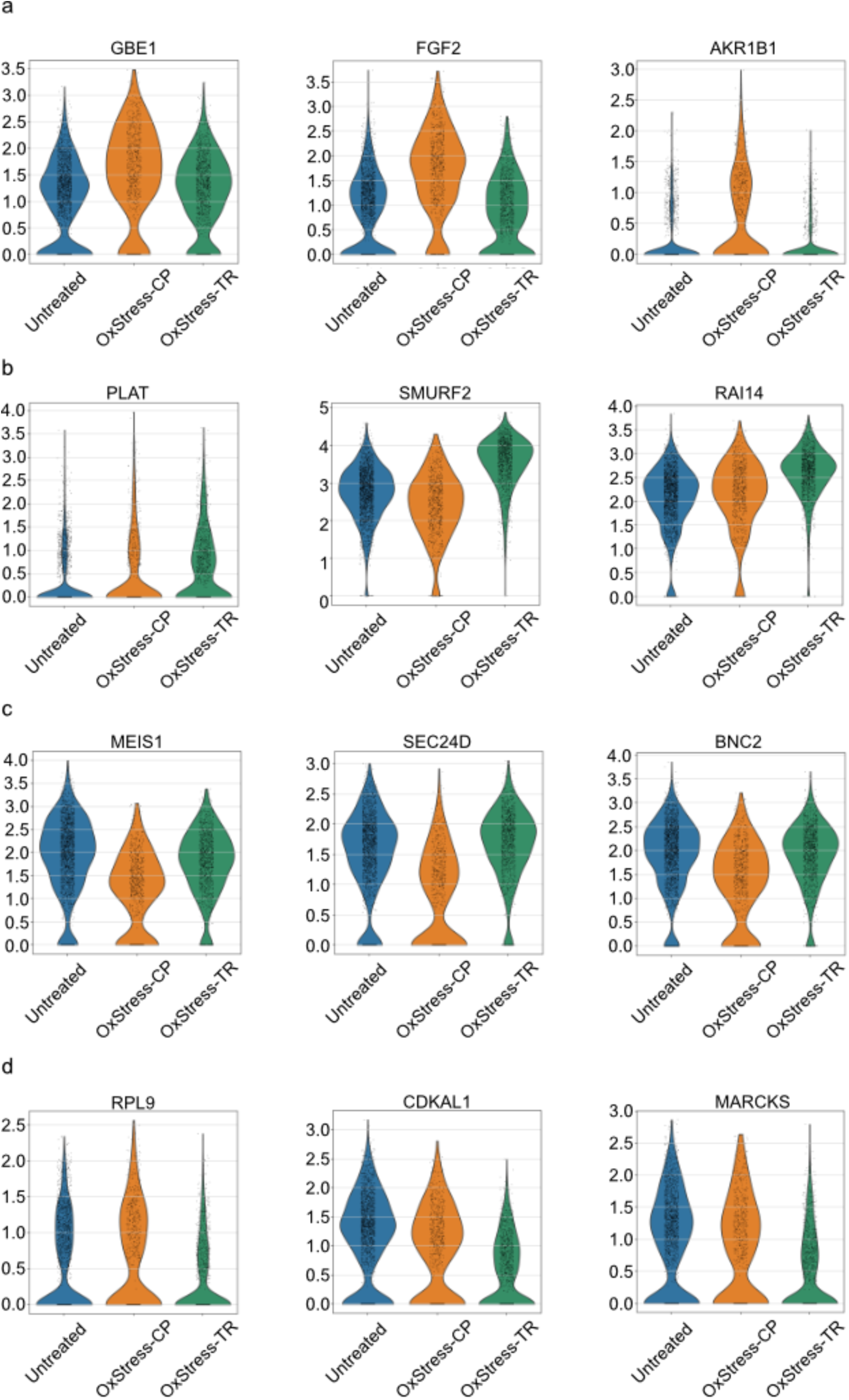
Subfractions of senescent cells differ in expression of some of senescence markers. **a**. Genes with previously reported increased expression in senescent cells with significantly higher expression in OxStress-CP compared to OxStress-TR. **b**. Genes with previously reported increased expression in senescent cells with significantly higher expression in OxStress-TR compared to OxStress-CP. **c**. Genes with previously reported decreased expression in senescent cells with significantly lower expression in OxStress-CP compared to OxStress-TR. **d**. Genes with previously reported decreased expression in senescent cells with significantly lower expression in OxStress-TR compared to OxStress-CP.

## Discussion

Recent studies highlighted growing variability of gene expression with age both *in vivo* and *in vitro* (Bahar et al., 2006; Cheung et al., 2018; Hernando-Herraez et al., 2019; Martinez-Jimenez et al., 2017; Salzer et al., 2018; Wiley et al., 2017). Given stochastic nature of molecular damage events, such as DNA damage, associated with stress and aging, such variability is to be expected and may give better insight into proximal causes of aging. Here, we used advances in genomic technologies which allowed scalable and multiplexed single-cell RNA-sequencing. We were able to perform multiple independent experiments which allowed us to verify reproducibility of the observed gene expression variation and perform its extensive characterization.

We examined cells induced into senescence with oxidative stress, since senescence was previously associated with increased gene expression variation as measured with single cell qPCR (Wiley et al., 2017). We extended prior findings with whole transcriptome analysis. We indeed found that induction of senescence through molecular damage is associated with increased cellular heterogeneity, as cells split into two transcriptional clusters. This was unlike cells induced into quiescence by low serum treatment or into senescence/senescence-like arrest with nutlin, which formed singular transcriptional clusters. Our data indicates that distinct cell fates start emerging early during stress response, potentially reflecting either pre-existing heterogeneity among untreated cells or stochastic differences in molecular damage during treatment. Metabolic difference emerged as a prominent feature that differentiate two post-stress fractions cells and it was supported by abundance of both mitochondrial and glycolytic transcripts. Given a central role of metabolic pathways in aging and stress adaptation (Fontana et al., 2010), it is tempting to speculate that difference in metabolic signaling may be a driving force for two distinct cells fates. Further investigations will allow to test this hypothesis and better understand the relationship between metabolism and senescence and potentially different subtypes of senescence.

Our results are consistent with the recent reports of two distinct senescent fates in the oncogene-induced senescence model (Hoare et al., 2016; Teo et al., 2019). Similarly, to those results we also observe two senescence fates, one of which is characterized by TGF-beta and Notch signaling. Here, we extended prior studies and performed a more detailed time course analysis of the emergence of senescence fates, and we found that bias toward a particular senescence fate emerges early during response. Prior reports labeled the observed fates as primary and secondary senescence with the idea that Notch signaling of primary senescent cells gives rise to secondary senescent cells. We are yet to see if similar hierarchy is applicable in our system, however, early emergence of fate bias suggest that other mechanisms also come into play. It is interesting to notice that nutlin-3a treatment didn’t give rise to the inflammatory fate, but only to the fate that was transcriptionally close to OxStressTR (see Figure 1). This is consistent with the previous report that nutlin-3a attenuates inflammatory phenotype of senescent cells (Wiley et al., 2018). It also suggests, that intensity and duration of p53 signaling may be another determining factor for the particular senescence fate (Purvis et al., 2012; Reyes et al., 2018; Tsabar et al., 2020).

In our experimental system, we did not find increased stochastic noise of gene expression. The cells that we measured were not more dissimilar from one another. These results are distinct from some previous reports, but not all (Kimmel et al., 2019). We do not yet understand the cause of these differences. It seems likely that different cell types and different senescence conditions will have different effects on the amount of cell-to-cell variation in gene expression and other kinds of cellular heterogeneity. It is suggested by Anderson et al that there will be both abnormal increases and decreases in cellular heterogeneity that will both have pathological consequences (Mendenhall et al., 2021). Only since the 21^st^ century have we begun to understand the causes and consequences of cell-to-cell variation in gene expression (Elowitz et al., 2002; Raser and O’Shea, 2004). Further in vivo and in vitro studies on cell-to-cell variation in senescent cells will likely yield additional means to target these cells for elimination to improve the health of aging animals and people. Our work suggests that the number of transcriptional states may be fairly limited among any particular cell type that becomes senescent, offering hope to the idea that a few different senolytic drugs may one day be sufficient to greatly improve human health.

Finally, various types of molecular stress induce formation of senescent cells with distinct transcriptional profiles. It will be important to analyze these different types of senescent cells at single-cell level to better understand the source of the differences. We may find that different stressors induce completely distinct types of senescent cells, or on the opposite, both shared and distinct subtypes of senescent cells may emerge after different types of molecular stress. Indeed, in our own system, we observed that one of the post-stress fractions is transcriptionally more similar to nutlin-treated cells, while another fraction is more distinct. It will be important to expand such single-cell analysis to different types of senescent cells for better understanding of the senescence and identification of more reliable markers of senescence for research and therapeutic applications.

## Methods

### Cell Culture and Treatments

Human fetal lung fibroblasts IMR-90 were cultured in Dulbecco’s modified Eagle’s medium containing 10% fetal bovine serum and penicillin/streptomycin. Cells were maintained at 37^°^C and 5% CO_2_. For treatments, cells were seeded at ~10^4^/cm^2^ and all treatments started next day. For induction of quiescence cells were kept in media with 0.2% serum for 7 days. For induction of senescence/senescence-like arrest with nutlin-3a, cells were kept for 7 days in media supplemented with 2.5uM nutlin-3a. For induction of senescence with oxidative stress, cells treated with 55uM hydrogen peroxide for two hours. In one experiment (Figure S1h-n), cells were treated with 75uM hydrogen peroxide twice, on Day 0 and Day 3. As in other experiments, cells were collected on Day 7 after the first treatment. In this experiment, quiescence was induced by low serum treatment for 3 days, from Day 4 till Day 7. In time course experiments, cells were collected at different time points after stress treatment, that was performed on Day 0. For assaying senescence-associated ß-galactosidase activity, cells were fixed with formaldehyde/glutaraldehyde and stained with 5-bromo4-chloro-3-indolyl P3-D-galactoside (X-Gal) as described before (Dimri et al, 1995).

### Preparation of cDNA library and sequencing

We followed standard SPLITseq protocol as described before (Rosenberg, Rocco et al). Briefly, cells were fixed in formaldehyde, then permeabilized with triton, filtered through 40um strainer and counted. Cells were subjected to 3 rounds of barcoding using 3’ targeting polyT oligos in the first round. After barcoding, cells were digested with proteinase K and isolated cDNA was further amplified, size-selected and indexed. The libraries were subjected to 2×150bp paired-end sequencing on Illumina platforms by Genewiz.

### Determining transcriptional profiles of individual cells

Reads were aligned to human genome (hg19) using STARSolo following developers guidelines. Read2 was used to decode cellular barcodes and unique molecular identifiers, while read 1 was actually mapped to transcriptome. Further processing was done with scanpy using developers guidelines (Wolf et al., 2018). We filtered out cells with low read count (less than 1000 or 2000 depending on sequencing depth) and cells with more than five percent of reads originating from mitochondrial genome. Genes expressed in less than 100 cells were excluded from analysis. We selected ~2000 most variable genes in each experiment and used them for principal component analysis (PCA). After PCA, cells were embedded using UMAP (Becht et al., 2019) and community detection was performed with Louvain algorithm (Blondel et al., 2008). Markers of transcriptional groups were determined by Wilcoxon rank-sum test.

### Gene set enrichment analysis (GSEA)

Gene set enrichment analysis was performed using its Python implementation, GSEAPY. Genes were ranked by Wilcoxon rank-sum-based comparison of different transcriptional groups. Ranked list of genes was used as an input for GSEAPY. We used ‘Hallmarks’ gene sets and performed the analysis with 1000 permutations of gene sets. Benjamini-Hochberg Correction was used for adjusted p values. For plotting, only gene sets with adjusted p values 0.05 or less and False Discovery Rate of 0.25 or less were included. When comparing two groups, positive enrichment scores indicate that gene sets are enriched in a first group compared to a second group, while negative enrichment scores indicate enrichment of gene sets in a second group compared to a first one.

### Scoring gene expression signatures

To derive gene expression signatures of senescent cells fractions, we selected genes that reproducibly show statistically significant (adjusted p values 0.05 or less) difference in expression between two fractions of senescent cells in independent experimental trials: in two non-time course experiments and three time course experiments. Final time point, Day 7, was used in time course experiments. Experiment with repetitive stimulation with hydrogen peroxide was not used to derive these gene expression signatures. Enrichment of the signatures in all cells of the time course experiments was performed using *score_genes* function of the *scanpy* Python library. To score enrichment of glycolytic genes, we used genes of the KEGG_GLYCOLYSIS_GLUCONEOGENESIS set.

## Acknowledgements

We thank Dr. Renuka Pillai for helpful edits and suggestions on the manuscript. We thank Drs. Anna Kuchina and Georg Seelig for technical consultation on single-cell RNAseq analysis.

